# An 8-week resistance training protocol is effective in adapting quadriceps but not patellar tendon shear modulus measured by Supersonic Shearwave Imaging

**DOI:** 10.1101/434423

**Authors:** P. Mannarino, T. T. Matta, F. O. Oliveira

## Abstract

Habitual loading and resistance training (RT) can determine changes in muscle and tendon morphology but also in its mechanical properties. Conventional ultrasound (US) evaluation of these mechanical properties present limitations that can now be overcome with the advent of Supersonic Shearwave Imaging (SSI). The objective of this study was to analyze the Vastus Lateralis (VL) and patellar tendon (PT) mechanical properties adaptations to an 8-week RT protocol using SSI. We submitted 15 untrained health young men to an 8-week RT directed knee extensor mechanism. VL and PT shear modulus (μ) was assessed pre and post intervention with SSI. VL muscle thickness (VL MT) and knee extension torque (KT) was also measure pre and post intervention to ensure the RT efficiency. Significant increases were observed in VL MT and KT (pre= 2.40 ± 0.40 cm and post= 2.63 ± 0.35 cm, p = 0.0111, and pre= 294.66 ± 73.98 Nm and post= 338.93 ± 76.39 Nm, p = 0.005, respectively). The 8-week RT was also effective in promoting VL μ adaptations (pre= 4.87 ± 1.38 kPa and post= 9.08.12 ± 1.86 kPa, p = 0.0105), but not in significantly affecting PT μ (pre= 78.85 ± 7.37 kPa and post= 66.41 ± 7.25 kPa, p = 0.1287). The present study showed that an 8-week resistance training protocol was effective in adapting VL μ but not PT μ. Further investigation should be conducted with special attention to longer interventions, to possible PT differential individual responsiviness and to the muscle-tendon resting state tension environment.

## 1 INTRODUCTION

Skeletal muscles act as active and primary motors for the body segments movements (1), while tendons represent an important connective tissue with high resistance to tensile loading, responsible for muscle force transmission to the bone (2). Both tendon tensile environment and muscle demand levels will determine adaptations in these structures (3,4). According to the overload environment, tendons can increase resistance and thickness or undergo inflammation and structural disorganization (5). Increased muscle demand and training regime, especially against an high external resistance, can result in hypertrophy, changes in architecture or, sometimes, lesions and degeneration (1). Due to its fundamental function for standing in upright position, walking and running and significant impairment in normal movement during pathologic situations, knee extensor mechanism (quadriceps and patellar tendon (PT)) mechanical properties has received particular interest recently (6–10).

In the last decades, changes in PT mechanical properties have been object of many studies. Its adaptations to overload regimen, resistance training (RT) protocols, ageing and pathologic conditions were extensively documented (6,11–13). Yet, the PT structural and histological adaptations to overloading are still not fully understood. Much of the knowledge about its mechanical changes *in vivo* is derived from indirect analysis based on B-mode ultrasound (US) imaging combined with dynamometry. This method, however, is subject to inaccuracies due to possible calibration errors and misinterpretations of the structures displacement (9,14). In the other hand, muscle mechanical properties analysis is not feasible using this strategy due the dynamic nature of muscle contraction, even in isometric actions, and the absence of intrinsic anatomical landmarks to calculate strain and displacement. Other methodologies, as an application of a durometer to describe a muscle hardness index are also very limited and the reproducibility of this specific device has not been systematically evaluated (15).

In that context, shearwave imaging technique received particular interest in US imaging routine, allowing a direct and static evaluation of tissues mechanical properties (8,16). Supersonic Shearwave Imaging (SSI) is able to determine directly the PT and quadriceps shear modulus (μ) in a determined region of interest (ROI) based on the combination of a radiation force and an ultrafast acquisition imaging system (17–19). The μ is therefore computed from the velocity of the propagating shear waves. SSI was already externally validated and PT μ showed a strong positive correlation to longitudinal Young’s Modulus (E), ultimate force to failure and resistance to tensile loading in *in vitro* models (20,21). Also, it has showed a strong positive correlation to the muscle loading environment passively or during active contraction (R^2^> 0.9) (22–24). SSI has also revealed PT μ reducing after stretch-shortening regimens, ageing, tendinopathy and surgical procedures (19,25–27). Analyzing muscle, SSI detected reductions in μ after long term stretching protocols (28), eccentric resistance training (29), aging (30), musculoskeletal pathologies (31) and μ increases during acute muscle damage (32).

Although PT mechanical adaptations and quadriceps structural changes after resistance training monitored by US are extensively documented (33,34), to our knowledge no previous studies evaluated the PT or quadriceps μ changes promoted by a RT protocol using SSI. Changes in tendon collagen cross-linking as well as muscle architecture can impact ultimate force to failure (20,21). These adaptations could be reflected in variations of tendon stiffness. Therefore, our initial hypothesis was that the PT stiffening and quadriceps hypertrophy after RT observed in previous studies using US evaluation, would be reflected as an increase in μ on SSI evaluation. Therefore, our main objective was to assess the changes in PT and quadriceps μ using SSI, after a progressive RT protocol.

## 2 MATERIALS AND METHODS

### 2.1 Ethics statement

The University Hospital Ethics Committee approved this study (registration number 2.811.595). The experimental procedures were conducted in accordance with the Declaration of Helsinki. All participants received instructions about the study procedures and provided informed written consent before testing.

### 2.2 Experimental procedure

The study was conducted at the Biomedical Engineering Department in our University. The body weight and height of all subjects were measured and body mass index (BMI) was calculated. Age and dominant leg were informed. All subjects’ PT and VL were submitted to SSI evaluation pre and post intervention. As a form to assure that the RT protocol was effective, Vastus Lateralis muscle thickness (VL MT) and knee extensor torque (KT) were measured at baseline and after the eight weeks of RT.

### 2.3 Subjects

In this longitudinal study, 15 untrained male volunteers (28.6 ± 3.26 years old, 177.3 ± 6.88 cm height and 91.8 ± 17.25 kg of body mass) had both knees examined. Age was set between 25 and 40 years old to eliminate any variation of PT properties due to age or gender. None of the subjects had participated in any systematic training or physical activity for at least 6 months. Any clinical history or report of knee pain/injuries, systemic disease or previous knee surgery was considered as exclusion criteria. All subjects were right handed and the right leg was used for analysis as a reference.

### 2.4 Resistance training protocol

Participants were designated to eight-week resistance training for the quadriceps femoris muscle consisting of free-weight Squats (SQ) and Knee Extensions (KE) in a knee extension machine (MatFitness^®^, São Paulo, Brazil) in this precise exercise order. RT protocol was designed based on the ACSM recommendations for healthy individuals and adapted based on previous studies with similar design (35). The frequency of the training program was 2 sessions per week with at least 72 hours rest between sessions. A total of 16 sessions were performed in the 8-week training period with all the sessions occurring between 8 and 10 AM.

At baseline, 10RM testing was performed for both exercises. All subjects were submitted to a familiarization before testing during which the subjects performed the same exercises as used in the 10RM tests with the aim of standardizing the technique of each exercise. The tests and retest were then performed on 2 nonconsecutive days separated by 48-72 hours. The heaviest resistance load achieved on either of the test days was considered the pre-training 10RM of a given exercise. The 10RM was determined in no more than five attempts, with a rest interval of five minutes between attempts and a 10-minute recovery period was allowed before the start of the testing of the next exercise.

The 10RM tests were used to set the initial training load. Subjects were instructed to perform both exercises to failure in all sets and the weighs were continually adjusted to keep the exercises in an 8-12 repetitions range, with a two-minute rest interval between sets. The RT program followed a linear periodization with progressive volume, according to the training schedule. Before each training session, the participants performed a specific warm-up, consisting of 20 repetitions at approximately 50% of the resistance used in the first exercise of the training session (SQ). Adherence to the program was superior to 90% in all individuals and a strength and conditioning professional and a physician supervised all the training sessions. Verbal encouragement was provided during all training sessions.

### 2.5 Measurement of patellar tendon shear modulus

An Aixplorer US (v.11, Supersonic Imaging, Aix-en-Provence, France) with a 60-mm linear-array transducer at 4–15 MHz frequency was used in this study. The transducer was positioned at the inferior pole of patella and aligned with the patellar tendon, with no pressure on top of a generous amount of coupling gel. B-mode was used to locate and align the PT longitudinally. When a clear image of the PT was captured, the shear wave elastography mode was then activated. The transducer was kept stationary for approximately 10 seconds during the acquisition of the SSI map. A total of four images were acquired and saved for off-line processing analysis. Scanning of PT was performed with the subject in supine lying and the knee at 30° of flexion (36). The knee was supported on a custom-made knee stabilizer to keep the leg in neutral alignment on the coronal and transverse planes (Fig 1). Prior to testing, the subjects were allowed to have a 10-min rest to ensure the mechanical properties of PT were evaluated at resting status. The room temperature was controlled at 20° C for all image acquisitions and the same experienced operator performed all exams.

**Fig. 1.**
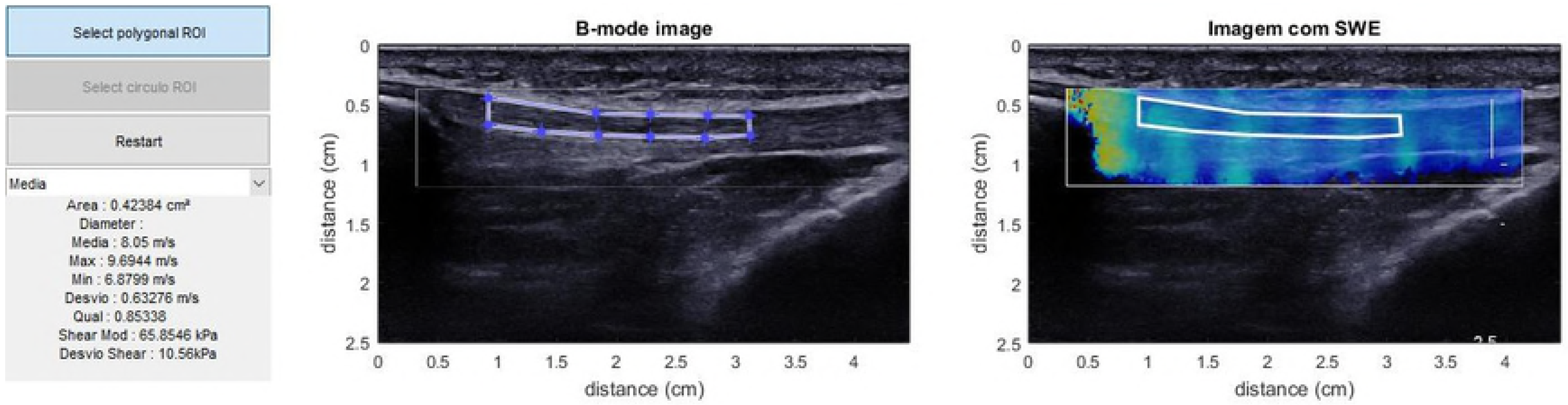
Imaging acquisition with the knee resting over a custom-made support at 30°.

The Q-box selected was the larger possible rectangle in order to consider more PT elasticity information. The μ values were obtained by a custom MatLab^®^ routine and ROI limits were defined as the area between 5 and 25 mm from the inferior pole of the patella excluding the paratendon (Fig 2) (37). The custom routine calculated the μ by dividing the mean E generated from the system by 3 (38).

**Fig. 2.**
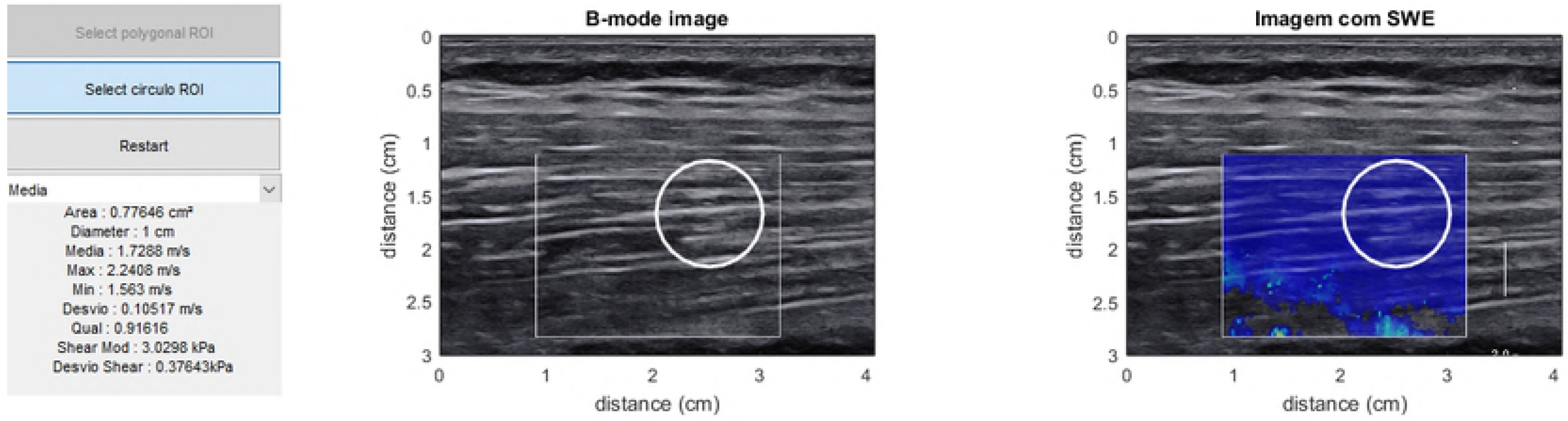
MatLab^®^ custom routine and ROI defined between 5 and 25 mm from the patella tip.

### 2.6 Measurement of Vastus Lateralis shear modulus

The same equipment with a 60-mm linear-array transducer at 4-15 MHz frequency was used for VL μ measurement. The US probe was centered and the images were recorded with subjects lying supine with their knee fully extended and their muscles relaxed. Scans were taken with a minimum compression and a generous amount of coupling gel at 50% of the length of the right thigh, represented by the distance from the great trochanter to the center of patella. The VL images were obtained on longitudinal plane laterally. B-mode was used to locate and align the VL longitudinally. When a clear image of the VL was captured, the shear wave elastography mode was then activated. The transducer was kept stationary for approximately 10 seconds until SSI map stabilization. A total of four images were acquired and saved for off-line analysis. Prior to testing, the subjects were allowed to have a 10-min rest to ensure the mechanical properties of VL were evaluated at resting status. The room temperature was controlled at 20°C. The same experienced operator performed all exams. The ROI was selected avoiding any detectable vascular structure within the muscle and based on the quality map (Fig 3). The custom routine calculated the μ by dividing the mean E generated from the system by 3 (38).

**Fig. 3.**
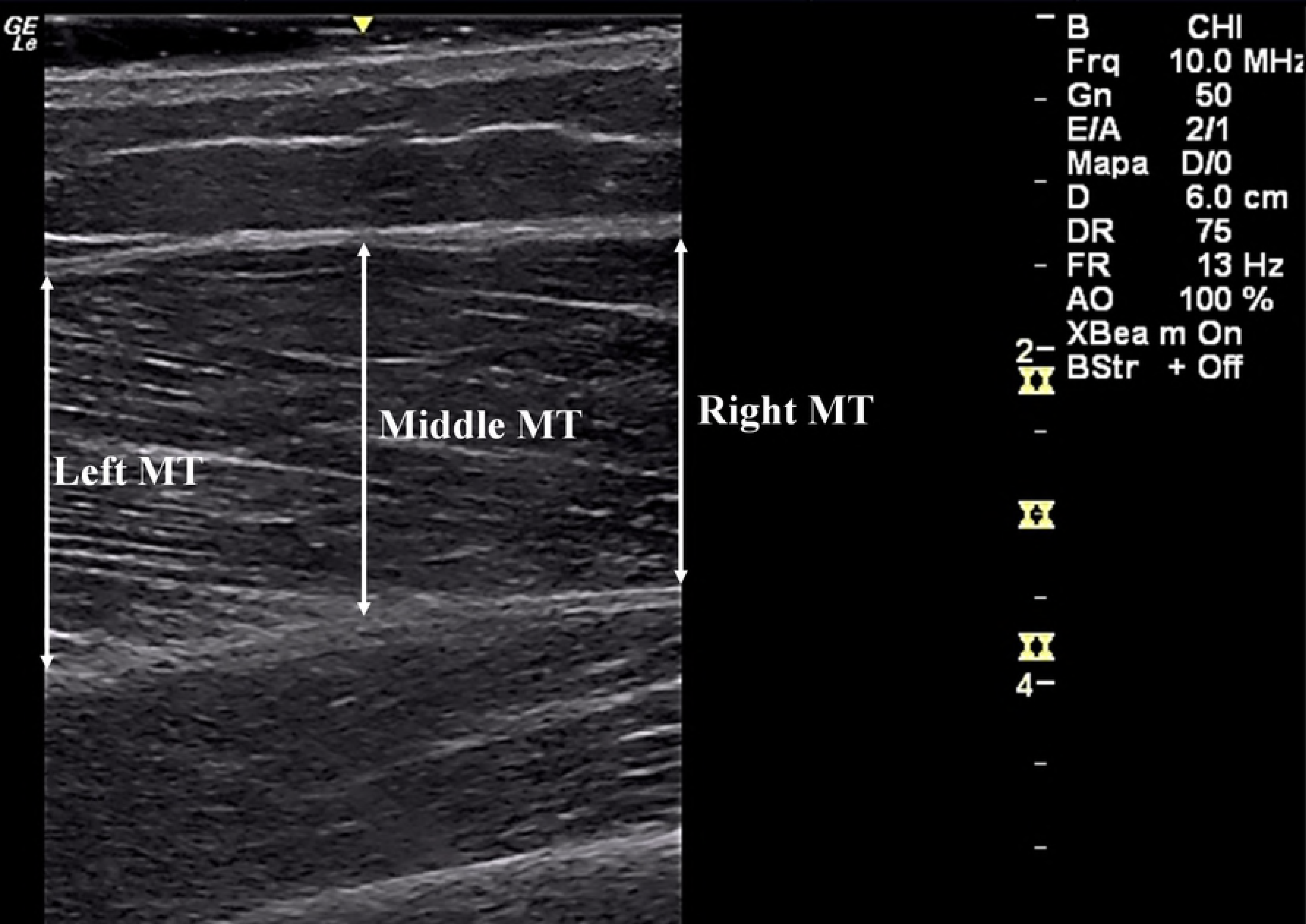
Vastus lateralis shear modulus measurement and selected ROI.

### 2.7 Measurement of Vastus Lateralis muscle thickness

The images acquisition was performed by an experienced researcher, using a US (GE LogiqE, Healthcare, EUA), frequency of 8 MHz, for longitudinal scans of the VL muscle. The US probe was centered and the images were recorded with subjects on the same position used for SSI. Scans were taken with a minimum compression a generous amount of coupling gel at 50% of the length of the right thigh, represented by the distance from the great trochanter to the center of patella. The VL images were obtained on longitudinal plane laterally and the MT was determined between the mean of three distances (proximal, middle and distal) between superficial and deep aponeurosis for each image **(39)**(Fig 4). The images were processed with publicly available software (ImageJ 1.43u; National Institutes of Health, Bethesda, MD, USA). For each image, two consecutive measurements were performed and the mean values were considered for analysis.

**Fig. 4.**
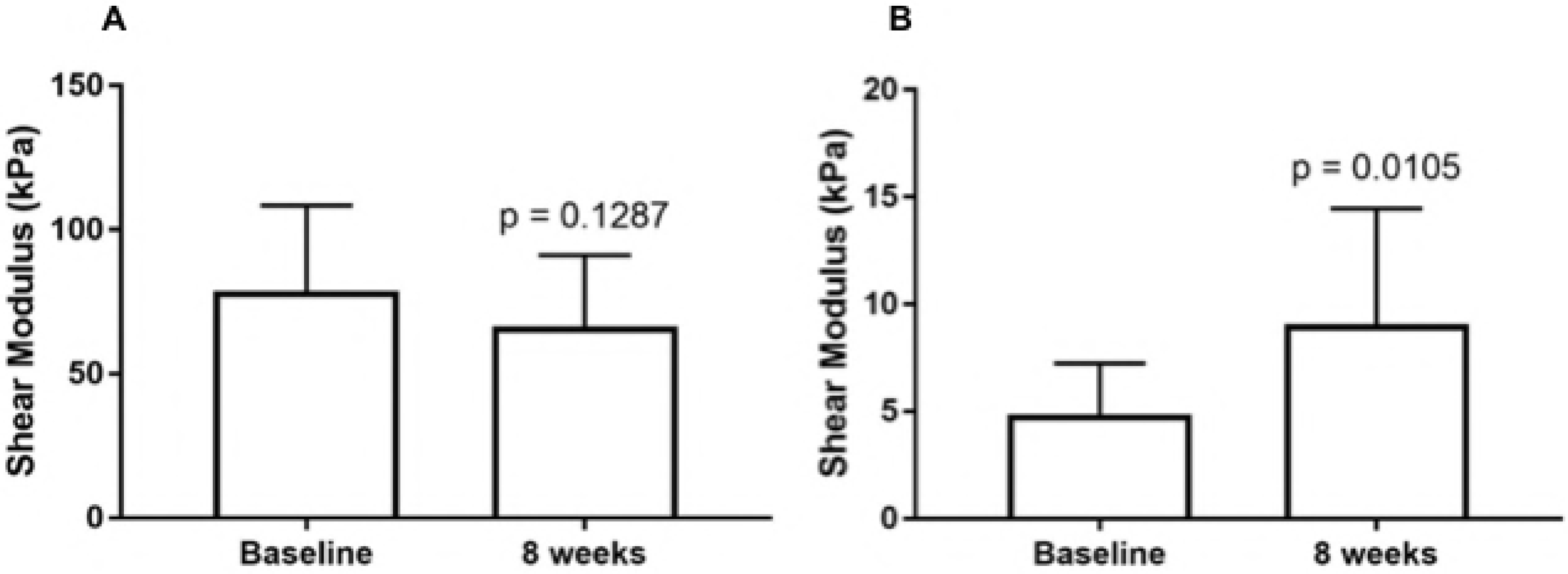
Vastus lateralis muscle thickness measurement with B-mode US.

### 2.8 Measurement of knee extension torque

The maximal isometric extension KT was measured with an isokinetic dynamometer (BIODEX^®^, Biodex Medical Systems, Shirley, NY, USA) at 80° of knee flexion (40). Subjects were positioned seated with inextensible straps fastened around the waist, trunk and distal part of the thigh. The backrest inclination and seat translation as well as the dynamometer height were adjusted for each subject, to ensure proper alignment of the rotation axis of the dynamometer with the lateral condyle of the femur. The right knee was fixed to the dynamometer lever arm 5 cm above the lateral malleolus. Settings were recorded for re-test reproducibility. After a specific warm-up consisting of two submaximal isometric knee extensions, the subjects performed two 5-s maximal voluntary isometric contractions (MVIC) with one-minute rest between trials. Subjects were verbally encouraged to reach maximal effort while a visual feedback of the torque level was provided. The highest peak torque among trials (corrected for gravity) was recorded for analysis.

### 2.9 Statistical analysis

Descriptive data such as mean ± standard deviation (SD) were calculated. The software GraphPad Prism 7^®^ was used for statistical analysis. After the normality distributions were assumed using the Shapiro-Wilk tests, paired t-tests were used to compare the PT μ, the VL μ, the VL MT and the KT at baseline and after the resistance training protocol. A value of p < 0.05 was adopted as statistically significant.

## 3 RESULTS

### 3.1 Patellar tendon and Vastus Lateralis shear modulus

No statistically significant changes in PT μ were observed after the eight weeks of RT (baseline= 78.85 ± 7.37 kPa and post = 66.41 ± 7.25 kPa, p = 0.1287). A statistically significant increase in VL μ was observed after the eight weeks of RT (baseline = 4.87 ± 1.38 kPa and post = 9.08.12 ± 1.86 kPa, p = 0.0105) (Fig 5).

**Fig. 5.**
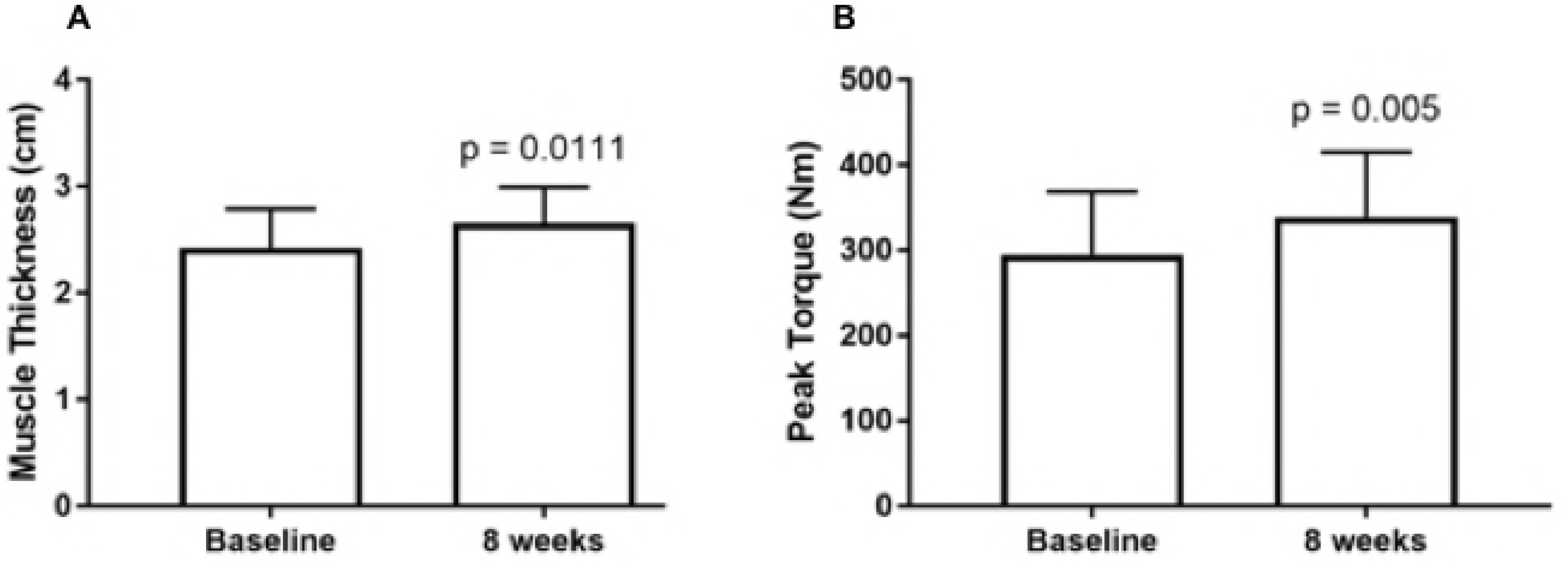
A- PT μ at baseline and after eight weeks of resistance training. B- VL μ at baseline and after eight weeks of resistance training.

### 3.3 Vastus lateralis muscle thickness and knee extensor torque

A statistically significant increase was observed in VL MT after the resistance training protocol (baseline = 2.40 ± 0.40 cm and post = 2.63 ± 0.35 cm, p = 0.0111). A statistically significant increase was observed in KT after the resistance training protocol (baseline = 294.66 ± 73.98 Nm and post = 338.93 ± 76.39 Nm, p = 0.005) (Fig 6).

**Fig. 6.**
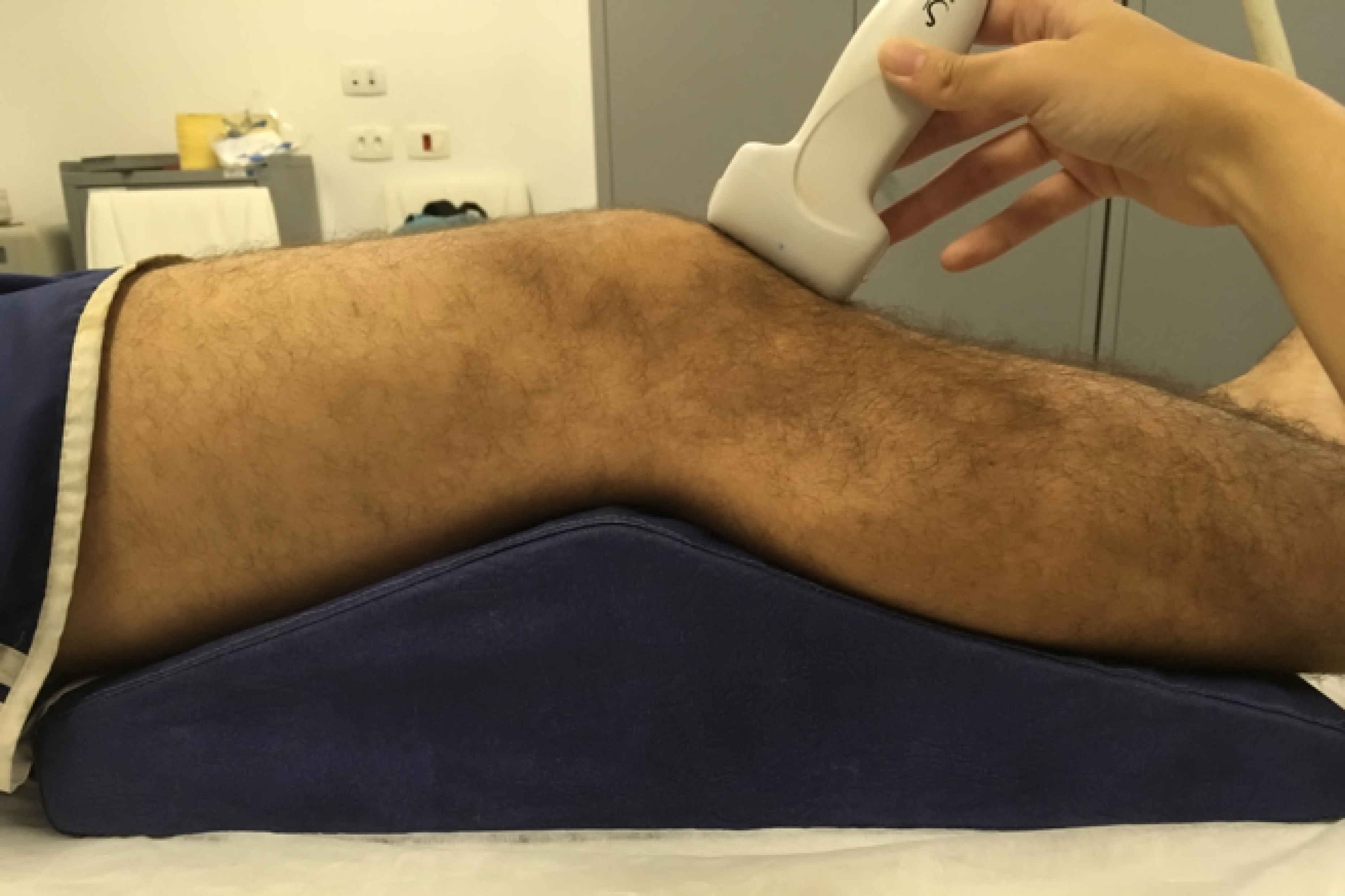
A- VL MT at baseline and after eight weeks of resistance training. B- KT at baseline and after eight weeks of resistance training.

## 4 DISCUSSION

Our study aimed to assess the effects of a progressive RT protocol directed to the knee extensor mechanism on the PT and VL μ measured by SSI. Our findings indicate that the proposed intervention was effective in promoting quadriceps hypertrophy and strength gains. VL MT and KT increased significantly (2.40 ± 0.40 cm at baseline and 2.63 ± 0.35 cm at 8 weeks, p = 0.0111, and 294.66 ± 73.98 Nm at baseline and 338.93 ± 76.39 Nm at 8 weeks, p = 0.005, respectively). Also, the intervention was effective in promoting VL μ adaptations (4.87 ± 1.38 kPa at baseline and 9.08.12 ± 1.86 kPa at 8 weeks, p = 0.0105), but not in significantly affecting PT μ (78.85 ± 7.37 kPa at baseline and 66.41 ± 7.25 kPa at 8 weeks, p = 0.1287).

### Effects of resistance training on patellar tendon shear modulus

The study of the tendons and muscle adaptation process to progressive overload is fundamental to design optimal strategies to injuries prevention and rehabilitation (41). Two recent literature reviews investigated the PT mechanical properties changes, with other techniques but SSI, after short-term (6-14 weeks) RT protocols (33,42). Significant increase in PT stiffness and E were observed routinely as a consequence of RT (10,43–45). In our study however, PT μ remained unaltered.

It is important to notice that these previous literature addressing the changes in PT mechanical properties secondary to RT were based exclusively on stress and strain estimation derived from B-mode US measures and dynamometry during isometric quadriceps contraction (33). Once there is no consensus related to the acquisition protocol, this methodology can jeopardize results previously found (9,14,46). Many technical details can limit these results reliability such as limited scanning from narrow fields of view (47,48), desynchronization between force production and elongation registration (49), limited three dimensional tracking of anatomical landmarks during muscle contraction (48,50,51), tendon force estimation inaccuracies (52) and others. Due to these practical limitations, we used SSI to observe the changes in PT shear modulus. To our knowledge, this is the first study analyzing PT μ changes after a RT protocol.

One relevant interpretation, is that the 8 weeks intervention could be not sufficient to trigger changes in tendon tissue composition at an extent to make the μ detectable by SSI. It has been previously reported that tendon remodeling process secondary to RT protocols can take longer periods (33), comparing to muscle adaptations. Although literature on the topic uses RT protocols lasting 6-14 weeks, a longer intervention could be necessary to make μ changes detectable by SSI. Our sample size and characteristics (15 individuals) was compatible to other studies previously published investigating PT adaptations to RT protocols with B-mode US (44,53), nevertheless, the known wide range of normal PT shear modulus values (7,8,37) may require a large sample to be tested. Future studies can, alternatively, apply classification techniques to identify responders and no responders individuals (54), as possibly the intervention could be effective for some of the individuals.

Lastly, the changes in resting state passive tension in the muscle-tendon unit deserves particular attention. It was previously reported that the shear modulus presents strong correlation to the tangent traction modulus at the time of SSI image acquisition (36). It is also documented that RT can increase flexibility (55) and that static stretching was able to reduce de E and μ acutely (56,57). This could mask the resistance training effects on PT μ. In one hand the expected increased collagen synthesis and tendon stiffening would increase shear modulus, while in the other hand the relaxation in the muscle-tendon unit and reduction in passive tension applied to the tendon could reduce it.

### Effects of resistance training on vastus lateralis shear modulus

Muscle mechanical properties seems to be much less explored, mainly by its limiting technical settings (58,59). Using US plus dynamometry technique seems not feasible due to the complex architecture when studying pennate muscles or due to the absence of reference anatomical landmarks in fusiform muscles to calculate strain. Until the advent of SSI, muscle mechanical properties were obtained by indirect analysis which is very limited, as the muscle hardness index using a durometer, whose reproducibility has not been systematically evaluated (15).

SSI values show a strong positive correlation with the muscle force production and activation, for example, for quadriceps, triceps surae, abductor minimum and others (22–24,60), showing that as muscle contract level rises, the more stiffer it becomes. However, the changes in muscle μ secondary to RT are far less studied.

To our knowledge, only two studies addressed this topic. Akagi et al. (2016) reported no changes in the triceps brachii μ after a 6-week RT consisting of triceps extensions (16). Differently from our study however, the authors report that transductor was positioned transversely to muscle fibers, which can actually exhibit lower μ values and blunt differences (61,62). Furthermore, the study used a shorter intervention (6 weeks) consisting of only one exercise in lower training volume (5 sets), what could explain the undetectable changes.

Another study investigated the effects of a 6-week protocol consisting exclusively of eccentric RT (Nordic Curl) on biceps femoris (29). Similarly, to Akagi et al. (2016) no statistically significant differences in biceps femoris μ were observed. However, in this study, stretching was also used in the protocol. It was already evidenced that stretching can reduce muscle μ (28), so increases in muscle μ secondary to the RT may have been nullified by the addition of a stretching intervention.

Increased stiffness in VL μ observed in our study could be resulted from collagen content, collagen linking and tissue fluid increasing (63). Also changes in muscle architecture and pennation angle as the VL, can have some impact in the μ, although the magnitude of these changes in the SSI technique is still under investigation (18,64).

### Limitations

Although SSI represents the state of the art in soft tissue mechanical properties evaluation (17,65), inferring structural adaptations in the musculoskeletal tissues after RT protocols is not trivial. Two different structures with similar composition can present different μ values on SSI evaluation if subjected to different tension during evaluation. The same applies to structures with different structural arrangement, that can present equal μ values on evaluation if subjected to different tension at the moment of testing (8,62,66). We controlled the testing position as much as possible to avoid this influence, trying to guarantee that the muscle was fully relaxed. Further studies researching muscle and tendon mechanical adaptations to RT with SSI should address this question and try to quantify the passive tension in the muscle-tendon unit in the resting state pre and post intervention.

To our knowledge, this is the first study analyzing the adaptation of the PT and VL mechanical properties to RT with SSI. Our initial hypothesis that RT would increase PT and VL μ was partially confirmed, once increases in VL μ were observed, but not in PT μ. These findings can help design further researches on the field and build new knowledge about the knee extensor mechanism remodeling process to mechanical overloading.

## 5 CONCLUSION

The present study showed that an 8-week resistance training protocol was effective in adapting VL μ but not PT μ, measured by SSI. Further investigation should be conducted with special attention to longer interventions, to possible PT structural changes set points and to the muscle-tendon resting state tension environment.

## ACKONWLEDGEMENTS

The authors acknowledge Financiadora de Estudos e Projetos - FINEP and Fundação de Amparo à Pesquisa do Estado do Rio de Janeiro - FAPERJ.

